# Addition of Daptomycin to Levofloxacin Increased the Efficacy of Levofloxacin Monotherapy against a Methicillin-Susceptible *Staphylococcus aureus* Strain in Experimental Meningitis and Prevented Development of Resistance

**DOI:** 10.1101/2020.10.28.360164

**Authors:** Philippe Cottagnoud, Frederike Sprenker, Marianne Cottagnoud, Alexandra Collaud, Reza Ashkbus, Vincent Perreten

## Abstract

Daptomycin and levofloxacin were tested as monotherapies and in combination against the antibiotic-susceptible *S. aureus* strain MSSA 1112 in a rabbit meningitis model and the effect of the combination on induction of resistance was determined in vitro. Changes of the susceptibility to fluoroquinolones and daptomycin was determined by the measurement of the MIC and mutations were detected by whole genome sequence comparison of the mutants with the parent strain MSSA 1112. Meningitis was induced by intracisternal inoculation of 10^5^ CFU of MSSA 1112 and treatment was started 10 h later by injection of daptomycin (15 mg/kg) and levofloxacin (10 mg/kg) standard doses. Cerebrospinal fluid (CSF) samples were repeatedly collected during therapy in order to determine killing rates and results of bactericidal activity were expressed in Δlog_10_ CFU/ml over 8 h. The combination of daptomycin with levofloxacin was significantly (p< 0.001) superior to levofloxacin monotherapy and increased the antibacterial activity of daptomycin. In vitro, MSSA 1112 was cycled over six days with either increasing concentrations of levofloxacin or daptomycin or with a combination of levofloxacin with half of the MIC of daptomycin or daptomycin with half of the MIC of levofloxacin leading to mutations in target genes as identified by whole genome sequence analysis. Addition of low concentration of daptomycin (0.25 mg/L) reduced levofloxacin-induced resistance in vitro. Addition of levofloxacin in low concentration (0.125 mg/L) did not influence daptomycin-induced resistance.

These findings highlight the lack of reciprocal interference of antibiotics in combination with regard to the development of resistance.

## Introduction

The treatment of meningitis caused by *Staphylococcus aureus*, occurring mainly after surgical interventions, remains a major challenge for clinicians and infectiologists (1). By now, the treatment of choice against methicillin-susceptible *S. aureus* (MSSA) strains is based on □-lactam antibiotics, and in case of penicillin allergy, vancomycin is usually used (2). Due to the adverse effects of vancomycin, i.e. nephrotoxicity and Red Neck syndrome, an alternative regimen with bactericidal activity is needed and would represent a considerable progress. Levofloxacin has been shown to have a good activity in pneumococcal meningitis (3) and daptomycin was efficacious as monotherapy in experimental meningitis caused by MSSA (4). However, development of resistance in *S. aureus* during monotherapies is of major concern(5, 6). Resistance of *S. aureus* to fluoroquinolones mainly occurs through mutations in the quinolone resistance-determining regions (QRDR) of the topoisomerase II (GyrA and GyrB) and IV (GrlA and GrlB) (7, 8). The mechanism of daptomycin resistance is still not well understood, but mutations and up-regulation of mostly cell wall related genes such as those encoding for phosphatidylglycerol lysyltransferase MprF, the two-component cell wall stress response associated regulator WalKR, the cardiolipin synthetase Cls2, the peptidoglycan synthesis associated proteins PgsA and MurA1/2 have been shown to be associated with daptomycin resistance in *S. aureus* (8–13). Additionally, mutations in the staphyloxanthin synthesis gene *crtM*, as well as deletion of the *rpsU* gene encoding for the 30S ribosomal protein S21, of the hypothetical lipoprotein Dsp1, and of the alkaline shock protein Asp23 also lead to decreased susceptibility to daptomycin (11, 14).

Addition of □-lactam antibiotics in subminimal inhibitory concentration (sub-MIC) has been shown to prevent the emergence of quinolone resistance in an experimental rabbit meningitis model using pneumococcal strains (3, 15). Such previous observations prompted us to analyze the efficacy of levofloxacin and daptomycin monotherapies and of a combination thereof against a methicillin-susceptible *S. aureus* strain using the same experimental meningitis model. Additionally, we evaluated the effect of the combination of levofloxacin and daptomycin in cycling experiments in vitro on the emergence of resistance against both antibiotics and searched for mutations by comparative analysis of whole genome sequence of the mutants with the parent strain.

## Material and Methods

### Rabbit Meningitis Model

Induction of meningitis with methicillin-susceptible *S. aureus* strain MSSA 1112 and CSF sampling were performed in Young New Zealand White rabbits as described previously (3, 16). Strain MSSA 1112 was initially isolated from the blood sample of a patient with endocarditis and was susceptible to levofloxacin (MIC 0.25 mg/L) and daptomycin (MIC 0.5 mg/L) (17). Animals were anesthetized by intramuscular injections for induction of meningitis with 10^5^ CFU of MSSA 1112 instilled in the cisterna magna and for CFU sampling from CSF. The experimental protocol was approved by the ethical committee of the Canton of Bern, Switzerland (Veterinäramt des Kantons Bern). The cisterna magna was punctured again for periodic CSF sampling before and 1, 2, 4, 5, 6, and 8 hours after initiation of therapy. Daptomycin (15 mg/kg) and levofloxacin (10 mg/kg) were administered by a peripheral ear vein injection once 14h after induction of meningitis.

Colony forming unit (CFU) of MSSA 1112 was determined by 10-fold serial dilutions of CSF samples plated on Muller-Hinton agar plates and incubated overnight at 37° C. Twenty microliters of undiluted samples were also plated to allow a limit of detectability of 50 CFU/ml). The dilutions of CSF were compared to exclude significant carryover effects during therapy. The antimicrobial activity of the antibiotics during the 8-h treatment was calculated by linear regression analysis and expressed as change in log_10_ CFU per milliliter per hour and as change of viable count over 8 h. A value of 1.7 (log_10_ of the limit of detectability) was assigned to the first sterile CSF sample, and a value of 0 was assigned to any following sterile CSF sample. The results were interpreted as means ± standard deviation and statistical significance was determined using the Newman-Keuls method (18).

### Selection of levofloxacin and daptomycin-resistant derivatives in vitro

The tendency of levofloxacin and daptomycin, alone or in combination, to select for resistant mutants of MSSA 1112 was tested in Mueller-Hinton broth. Inocula of 10^7^-10^8^ CFU/ml MSSA 1112 were exposed to serial two-fold increasing concentrations of levofloxacin or daptomycin. After 12 hours of incubation at 37°C, 0.1 ml of the cultures from the tubes containing the highest antibiotic concentration and still showing turbidity were used to inoculate a new series of tubes containing antibiotic serial dilutions. The experiments were performed over 12 cycles, and the minimal inhibitory concentration (MIC) was determined at the end of the experimental period.

The same method was used either with daptomycin added in low concentration (0.25 mg/L corresponding to half the MIC (1/2 MIC) to the tubes containing two-fold serial dilutions of levofloxacin or with levofloxacin (1/2 MIC, 0.125 mg/L) to the tubes containing two-fold serial dilutions of daptomycin. The experiments were performed during 12 cycles.

### Antimicrobial susceptibility testing of the parent and mutant strains

Antimicrobial susceptibility was determined for *S. aureus* MSSA 1112 and derived mutants (MSSA 1112-2, MSSA 1112-3, MSSA 1112-4 and MSSA 1112-5) by the measurement of the MIC of ampicillin, ciprofloxacin, daptomycin, erythromycin, tetracycline, gentamicin, linezolid, quinopristin-dalfopristin, teicoplanin, tigecycline, and vancomycin in Mueller-Hinton broth using sensititre plates EUVENC Sensititre® plates (Thermo Fisher Scientific, Waltham, USA) and home-made 96-well microtiter plates for levofloxacin following the CLSI guidelines (19).

### Whole genome sequencing and detection of mutations associated with increased resistance to antibiotics

Whole genome sequencing (WGS) of *S. aureus* strain MSSA 1112 was performed with both long and short reads to obtain a complete and accurate genome to be used as scaffold. WGS of the 4 MSSA 1112 mutants (MSSA 1112-2 to MSSA 1112-5) was made with short reads which were used for read mapping against the scaffold of MSSA 1112. Short reads were obtained using NovaSeq6000 S2 Reagent Kit (300cycles) and S2 flow cell on a NovaSeq6000 (2×150-bp paired-end) system (Illumina, USA) at Eurofins Genomics GmbH, Germany. The adaptors and low-quality reads (QV ≤ 20) of the short reads were removed using Trimmomatic v0.36 (20). Long read sequencing of MSSA 1112 was performed using Oxford Nanopore Technologies (ONT) (ONT, United Kingdom). The library was prepared using the Ligation Sequencing Kit 1D SQK-LSK108 in an R9.4 SpotON flow cell with a MinION Mk1B device (ONT). The reads were base-called using Guppy basecaller (v2.3.7) (ONT). Hybrid *de novo* assembly of both short and long reads of MSSA 1112 was obtained using Unicycler v0.4.4 (21), and the resulting assembled genome was annotated using the NCBI Prokaryotic Genome Annotation Pipeline (22). Detection of mutations and read mapping of the Illumina reads of the 4 mutants (MSSA 1112-2 to MSSA 1112-5) against the genome of the parent strain MSSA 1112 was performed using with Geneious Prime® 2019.0.4 (Biomatters Ltd, Auckland, New Zealand). DNA sequence of genes containing mutations was translated into amino acid sequences to identify non-silent mutations at the protein level.

## RESULTS

The efficacies of the different antimicrobial regimens in experimental rabbit meningitis are presented in Table 1. At the beginning of the experimental period, the inocula were similar in all treatment groups. As described previously, untreated controls grew slowly and showed only a slight increase of the CFU after 8 hours (+ 0.60 ± 0.43 log_10_ CFU/ml) (4). Daptomycin monotherapy was bactericidal and showed a decrease of − 0.48 ± 0.13 log_10_ CFU/ml per hour and a decrease of − 4.06 ± 1.23 log_10_ CFU/ml after 8 hours. Levofloxacin monotherapy was slightly less efficacious with killing rates of 0.38 ± 0.05 log_10_ CFU/ml per hour and 3.47 ± 0.55 log_10_ CFU/ml at the end of the experimental period. Combination of both antibiotics was the most potent regimen with killing rates of 0.87 ± 0.09 log_10_ CFU/ml per hour and 4.81 ± 0.69 log_10_ CFU/ml after 8 hours, respectively. At the end of the experimental period, levofloxacin alone was not sufficient to clear the CSF of the rabbits from bacterial cells. Two of 5 CSF were sterile in the daptomycin group and 4 of 5 CSF in the combination group, respectively.

**Table 1.** Efficacy of the different antibiotic regimens against methicillin-susceptible *S. aureus* in experimental meningitis

| Groups (N)<br>[Strain] | Inoculum<br>(log<sub>10</sub> CFU/ml) | Killing rates/h<br>(Δ log<sub>10</sub> CFU/ml·h) | Killing rates/8h<br>(Δ log<sub>10</sub> CFU/ml·8h) |
| --- | --- | --- | --- |
| Controls (5)<br>[MSSA] | 5.53 ± 0.39 | + 0.08 ± 0.06 | + 0.60 ± 0.43 |
| DAP (5)<br>[MSSA] | 5.41 ± 0.37 | - 0.48 ± 0.13<sup>1,2</sup> | - 4.06 ± 1.23<sup>5</sup> |
| LVX (5)<br>[MSSA] | 5.54 ± 0.33 | - 0.38 ± 0.05<sup>1,3</sup> | - 3.47 ± 0.55<sup>4</sup> |
| LVX+DAP<br>(5)<br>[MSSA] | 5.21 ± 0.46 | - 0.87 ± 0.09<sup>2,3</sup> | - 4.81 ± 0.69<sup>4,5</sup> |

**1:** LVX versus DAP: p<0.17, not significant
**2:** LVX+DAP versus D: p=0.08, not significant trend
**3:** LVX+DAP versus LVX: p<0.001, highly significant
**4:** LVX+DAP versus LVX: p=0.009, significant
**5:** LVX+DAP versus DAP: p=0.26, not significant

Six days of cycling with levofloxacin alone led to an eightfold increase of the MIC (2 mg/L). The addition of daptomycin in subinhibitory concentration (0. 25 mg/L corresponding to 1/2 MIC) to levofloxacin cultures drastically reduced the increase of the MIC of levofloxacin (0.5 mg/l). Also after six days of cycling with increasing concentrations of daptomycin alone, the MIC of daptomycin increased to 4 mg/L. In contrast, the addition of low dose of levofloxacin (0.125 mg/L corresponding to 1/2 MIC) to daptomycin cultures did not influence the increase of the MIC of daptomycin (8 mg/L) (Table 2). In addition, the MIC for vancomycin was determined for all groups. Interestingly, cycling MSSA 1112 cultures with daptomycin led to a slightly increase of the MIC of vancomycin (MIC 2 mg/L). Even the addition of daptomycin in low dose to levofloxacin induced the same increase of MIC of vancomycin. On the other hand, levofloxacin monotherapy did not influence the MIC of vancomycin (Table 2). No change in the MIC of the other antibiotics tested was observed.

**Table 2.**
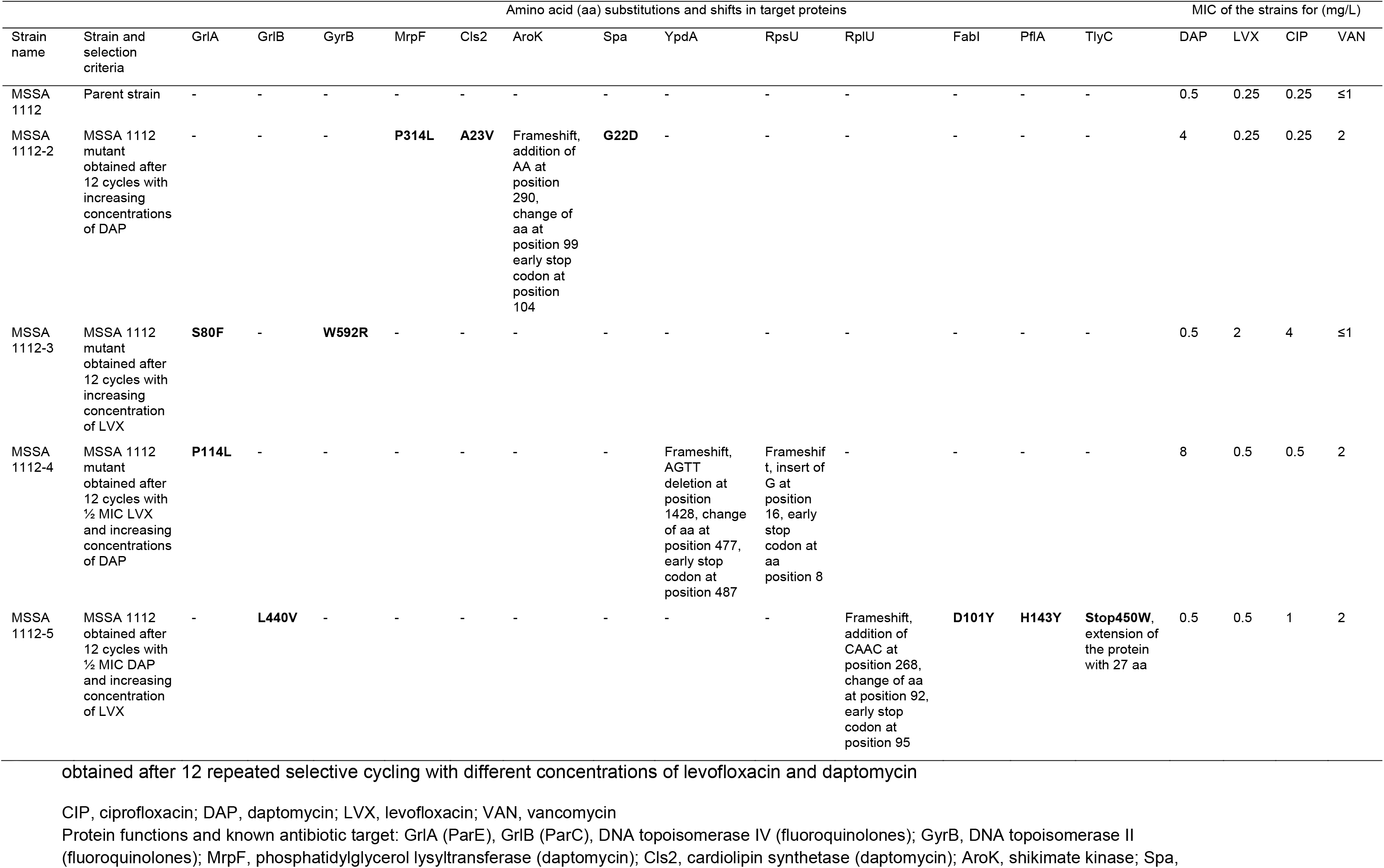
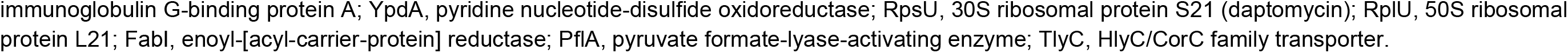
Amino acid substitutions in proteins and resulting minimal inhibitory concentration (MIC) changes of *S. aureus* MSSA 1112 mutants

The genome of the parent strain MSSA 1112 and the mutants was sequenced to identify mutations resulting in the mutants after the different antibiotic cycles (Table 2). Mutations associated with fluoroquinolone resistance were found in the QRDR of the topoisomerases IV (GrlA) and II (GyrB) of MSSA 1112-3 after cycling of MSSA 1112 with levofloxacin monotherapy. In mutant MSSA 1112-5 generated in the cycling group containing increasing concentrations of levofloxacin combined with a low dose of daptomycin, mutations resulted in amino acid substitutions in the QRDR of GrlB with slight increase of the levofloxacin MIC to 0.5 mg/L, and to amino acid substitutions in FabI, as well as frameshift in RpIU and TlyC resulting in an early stop codon and an extension of the protein sequence, respectively. None of these mutations affected the MIC of daptomycin which remained the same as the parent strain (Table 2). The two mutants MSSA 1112-2 and MSSA 1112-4 exhibiting decreased susceptibility to daptomycin harbored several different mutations. Mutant MSSA 1112-2 which was selected on daptomycin only harbored amino acid substitutions in MrpF, Cls2 and Spa, as well as frameshift-resulting mutations in AroK. Mutant MSSA 1112-4 which was selected with increasing concentration of daptomycin combined with low dose levofloxacin, exhibited an amino acid substitution in GrlA and two frameshift-resulting mutations in the *ypdA* and *rpsU* genes, both leading to an early stop codon in the respective proteins YpdA and RpsU (Table 2).

## Discussion

In previous studies, we have shown that □□lactam antibiotics, i.e. ceftriaxone, cefotaxime, and meropenem were very active especially in combination with levofloxacin in experimental pneumococcal meningitis (3, 15, 23). These studies also showed that addition of □-lactam antibiotics reduced the emergence of levofloxacin-induced resistance in pneumococci demonstrated in cycling experiments in vitro. In the present study, a similar experimental setting was used to test the efficacy of daptomycin combined with levofloxacin compared to the monotherapies against a MSSA strain in experimental meningitis. Furthermore, the potency of the combination to modify the emergence of resistance compared to the monotherapies was evaluated in vitro. As expected from previous experiments (4), bacterial growth was slow in the untreated controls over 8 hours. Daptomycin was slightly more efficacious than levofloxacin although the difference was statistically not significant. Only the combination treatment was clearly superior to the monotherapies, especially to levofloxacin (P<0.001). Probably a higher number of animals in each group would have been more discriminating, especially between the two monotherapies. Nevertheless, daptomycin as monotherapy or in combination with quinolones, seems to be a promising therapy of meningitis caused by MSSA.

In cycling experiments in vitro, incubation of the MSSA 1112 strain either with levofloxacin or daptomycin led to a pronounced increase of the MIC for the two monotherapies. The MIC increases were confirmed by detection of mutations in the target genes. Levofloxacin induced resistance was confirmed by the presence of a S80F substitution in GrlA, which is very common among fluoroquinolone-resistant clinical *S. aureus* isolates (7). An additional amino acid substitution was detected in GyrB (W592F) and its role in increased MIC of levofloxacin is not known. The mutant selected under daptomycin contained four different mutations, including amino acid substitutions in MprF (P314L) and Cls2 (A23V), which correspond to known mutations associated with resistance to daptomycin resistance in clinical *S. aureus* isolates and in in vitro generated mutants (9, 10, 24). The two other mutations were detected in the shikimate kinase AroK and the immunoglobulin G-binding protein A Spa, whose associations with decreased susceptibility to daptomycin are not known. The most striking feature of the study was the effect of the combination treatments on the emergence of resistance to the monotherapies. Addition of low-dose daptomycin to cultures cycled with levofloxacin reduced the development of resistance to levofloxacin leading only to a twofold increase of MIC of levofloxacin. In this mutant (MSSA 1112-5) only one amino acid substitution was detected in GrlB (L440V) which is situated within the QRDR of GrlB and may explain this slight increase of MIC (25). This mutant obtained in the presence of low dose of daptomycin harbored additional mutations, i.e. frameshift mutations in RplU, FabI, PflA and TlyC genes without influencing the MIC for daptomycin. Obviously these mutations are of minor relevance in creating daptomycin resistance. On the other hand, addition of low dose levofloxacin to cultures cycled with daptomycin did not influence the daptomycin-induced resistance. In this mutant, frameshift mutations were detected in pyridine nucleotide-disulfide oxidoreductase YpdA and the 30S ribosomal protein S21 (RpsU). While deletion of RpsU is known to confer resistance to daptomycin in *S. aureus* (14), the association of a frameshift mutation in YpdA with daptomycin resistance is so far not known. In this mutant, a mutation in GrlA was also induced by low dose levofloxacin, correlating with the slight increase of its MIC.

Another interesting feature was the effect of daptomycin selection on the slight increase of the MIC of vancomycin. The underlying mechanism of this resistance development affecting these two antibiotics with a different bacterial target (cell wall synthesis versus integrity of cell membrane) is not well understood. However, simultaneous decreased susceptibility of *S. aureus* to vancomycin and daptomycin during either vancomycin or daptomycin treatment has been reported in clinical isolates (26–29) (REF) (REF). Specific mutations in MprF have been shown to affect non-susceptibility to both daptomycin and vancomycin (30, 31). However, the MprF mutation (P314L) generated in the MSSA 1112-2 mutant has so far only been shown to be associated with non-susceptibility to daptomycin, both not to vancomycin (9). Additionally, the role of the remaining mutations in Cls2, AroK, Spa, YpdA, RpsU, RplU, PflA, FabI, TlyC generated under daptomycin selection in vancomycin non-susceptibility still remains to be determined.

In conclusion, daptomycin combined with levofloxacin seemed to be an efficacious and promising treatment for bacterial meningitis due to MSSA. In vitro, daptomycin prevented or decreased levofloxacin-induced resistance. On the other side, levofloxacin did not manage to influence daptomycin-induced resistance. Of note, both antibiotics added in low dose in the combination groups were able to induce mutations in their target genes emphasizing the importance of prudent and appropriate use of antibiotics.

## ACKNOWLEDGEMENTS

We thank Dr. J. Entenza and Dr. P. Moreillon of the Department of Internal Medicine, Centre Hospitalier Universitaire Vaudois (CHUV), Lausanne, Switzerland, who kindly provided *S. aureus* strain MSSA 1112.

